# Ultrasound exposure enhances cellular uptake of nanoscale cargos

**DOI:** 10.1101/2021.09.11.459898

**Authors:** Kahkashan Bansal, Anjali Rajwar, Himanshu Shekhar, Dhiraj Bhatia

## Abstract

DNA nanotechnology utilizes DNA as a structural molecule to design palette of nanostructures with different shapes and sizes. DNA nanocages have demonstrated significant potential for drug delivery. Therefore, enhancing the delivery of DNA nanocages into cells can improve their efficacy as drug delivery agents. Numerous studies have reported the effects of ultrasound for enhancing drug delivery across biological barriers. The mechanical bioeffects caused by cell-ultrasound interaction can cause sonoporation, leading to enhanced uptake of drugs, nanoparticles, and chemotherapeutic agents through membranes. Whether ultrasound exposure can enhance the delivery of DNA nanocages has not been explored, which is the focus of this study. Specifically, we investigated the effects of ultrasound on the cellular uptake of propidium Iodide, fluorescent dextrans, and DNA nanostructures). We provide evidence of modulation of pore formation in the cell membrane by ultrasound by studying the intracellular uptake of the impermeable dye, propidium iodide. Treatment of cells with low amplitudes of ultrasound enhanced the uptake of different sizes of dextrans and DNA based nanodevices. These findings could serve as the foundation for further development ultrasound-enabled DNA nanostructure delivery and for specific understanding of underlying biological mechanisms of interaction between ultrasound parameters and cellular components; the knowledge that can be further explored for potential biological and biomedical applications.

## Introduction

Ultrasound imaging employs non-ionizing waves that can penetrate several cm in the body and provide diagnostic information. Ultrasound also has thermal and mechanical effects that can be used for therapeutic applications. Therapeutic ultrasound frequencies typically range from 200 kHz to 2 MHz. Many researchers have utilized ultrasound for the delivery of molecules, nanoparticles, chemotherapeutic agents to name a few.

DNA nanotechnology has been extensively applied for numerous biological and biomedical applications. DNA-based nanostructures have gained enormous attention in the scientific research owing to their ability to be coupled to external biological targeting entities like peptides, small molecules, antibodies, etc., and their ability to encapsulate various nanoscale cargo like quantum dots, drugs, small molecules etc. within their internal void. Because of its programmability, inherent biocompatibility, chemical flexibility, DNA serves as a next-generation biomaterial for therapeutics development. Compared to other polymer or protein-based nanomaterial, DNA-based scaffolds provide precise control over the spatial arrangement of different moieties and have been efficiently utilized as delivery vehicle. However there are some roadblocks for utilizing the full potential of these DNA nanodevices like efficient cellular delivery of DNA nanodevices being one of them. Multiple methods have been employed for enhancing the uptake of DNA nanodevices into biological systems like coupling them with proteins, lipids, sugars, or transfection reagents, etc. However, not much has been explored on modulating the surface properties of cells and tissues using biophysical methods and studying its effects on DNA nanodevices uptake into cells.

We herein studied the US-mediated delivery of DNA nanocages (NCs) by using various US parameters [Karimi M, 2017]. To optimize ultrasound parameters, studies were first conducted using PI and FITC-Dextran. To test whether sonoporation affects cell viability, we assessed whether sub-cellular organelles remain intact and unstressed. Using various markers for endocytosis and cellular organelles, we provide the evidence of modulation of pore formation in membrane by ultrasound by studying the intracellular uptake of the impermeable dye, Propidium iodide. Treatment of cells with low intensity ultrasound enhanced the uptake of endocytic markers like fluorescent dextrans and DNA based nanodevices, specifically 3D DNA nanocages. We also found that the morphology of cellular components such as mitochondria, lipid membranes, neutral lipid molecules, actin filaments remain unchanged after the treatment, indicating the safety of ultrasound for application to cells for the range of exposure parameters employed.

These findings could serve as the foundation for further development ultrasound-enabled DNA nanostructure delivery and for specific understanding of underlying biological mechanisms of interaction between ultrasound parameters and cellular components.

## Results

### 1. Effect of ultrasound on uptake of cell-impermeable dyes

The influence of ultrasound (US) intensity on cells was evaluated by monitoring the uptake pattern of propidium iodide (PI), a fluorescent membrane impermeable compound. The US treatment was given to cells seeded on 35mm dish by placing the dish on the top of the probe with ultrasound gel. SUM159 triple negative breast cancer cells were exposed to ultrasound treatment of varying intensities with the exposure time of 1 min. After treatment cells were incubated with 1µg/ml PI solution in Ham’s F12 complete media for 10mins. Non-treated cells were considered as control and were incubated with PI for 10 mins without any ultrasound exposure. The PI intensity showed a marked increase with US exposure at 5.6 and 7 W/cm^2^ intensities **(Fig 1a, b)**. We varied the exposure time from 0 sec to 120 sec while keeping the intensity at 5.6W/cm^2^. We observed that the internalization of PI was maximum at 60 sec **(Fig 1c, d)**. However, 7 W/cm^2^ exposure intensity for 60 sec, caused the detachment of cells from the cell well plate and changes were observed in their morphology. Nonetheless, these studies indicate that sonoporation was occurring in the cells as PI, a membrane impermeable dye, was able to penetrate the cells.

**Figure 1.**
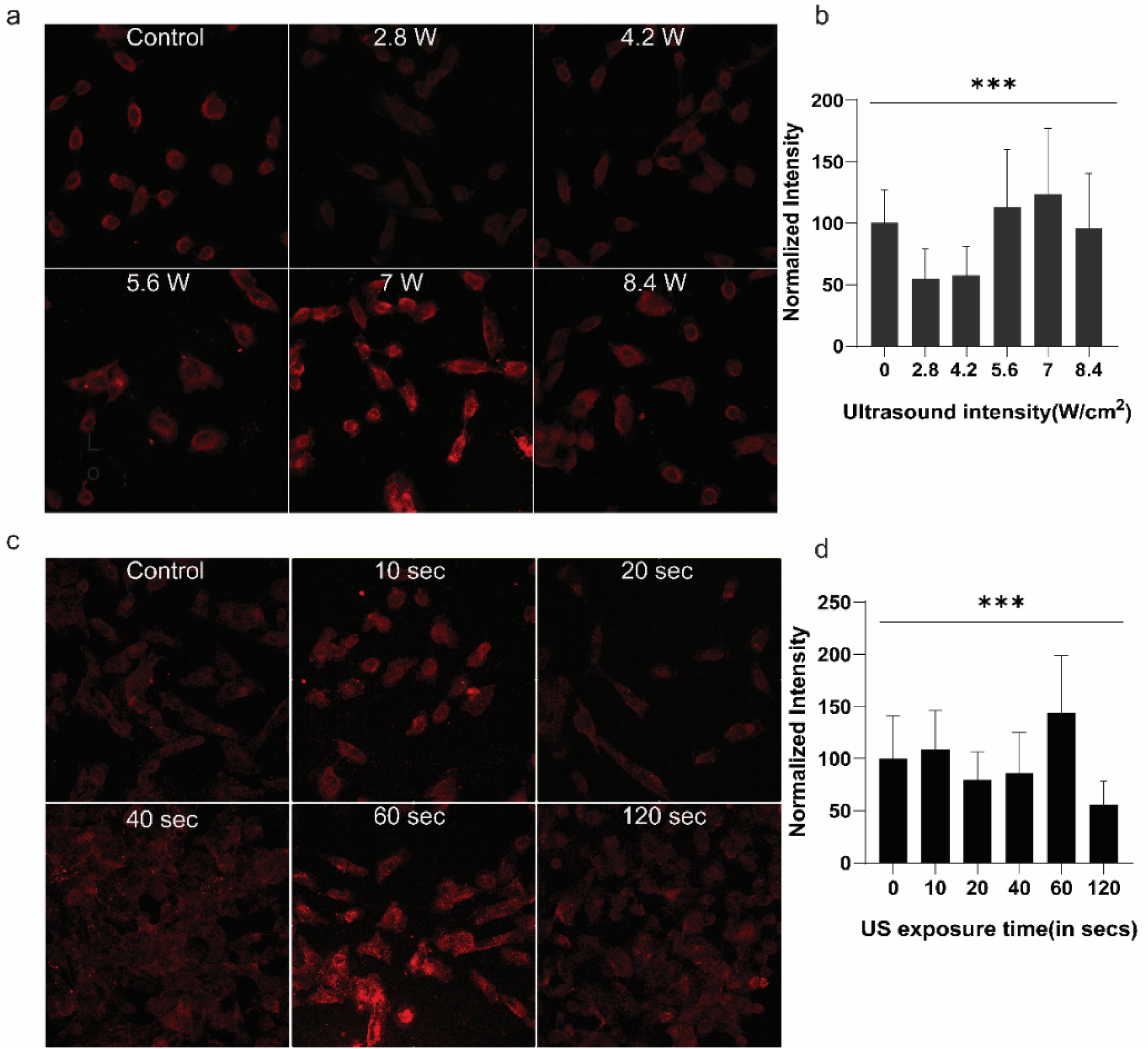
Influence of ultrasound on PI internalization. **(a)** Confocal images of SUM159 cells showing the internalization of PI upon exposure to different acoustic intensities. Images were taken in z-stacks and analyzed using FIJI ImageJ. **(b)** Quantification of PI penetration upon US treatment from cells in panel (a). **(c)** Penetration of PI in cells at different time points with anUS exposure of 5.6 W/cm^2^. **(d)** Quantification of PI permeation at different time-points from cells in panel (c). The internalization of PI changes with exposure time. Error bar indicates mean ± SD of PI intensity in each condition. Normalized intensity was calculated from > 30 cells per condition. *** indicates p value <0.0001.

We compared the intensities between serum and serum-free samples to assess its effect on PI delivery. These were performed for two PI concentrations-1 µg/ml and 2 µg/ml **(Figure 2)**. The US intensity and US exposure time were both kept constant-7 W/cm^2^ and 10sec respectively. The incubation time with PI after US exposure was kept constant for 10 min.” For 1 µg/ml concentration of PI, the control cells in complete media showed autofluorescence whereas no autofluorescence was detected in serum free media under same imaging conditions. However, in 2 µg/ml PI concentration the autofluorescence was detected in both complete and serum free media **(Figure 2a)**. We observed the concentration dependent internalization of PI showing more fluorescence signal at 2 µg/ml compared to 1 µg/ml under same imaging conditions **(Figure 2a)**. The cells in complete media showed no significant difference in fluorescence intensity upon ultrasound treatment in both PI concentration **(Figure 2b(i))**. However, the US treatment of cells in serum free media showed significant difference in the signal of treated and control cells in both PI concentrations **(Figure 2b(ii))**. This suggested that the DNase present in the serum degrades the DNA, so PI was unable to intercalate properly resulting in lower signal.]

**Figure 2.**
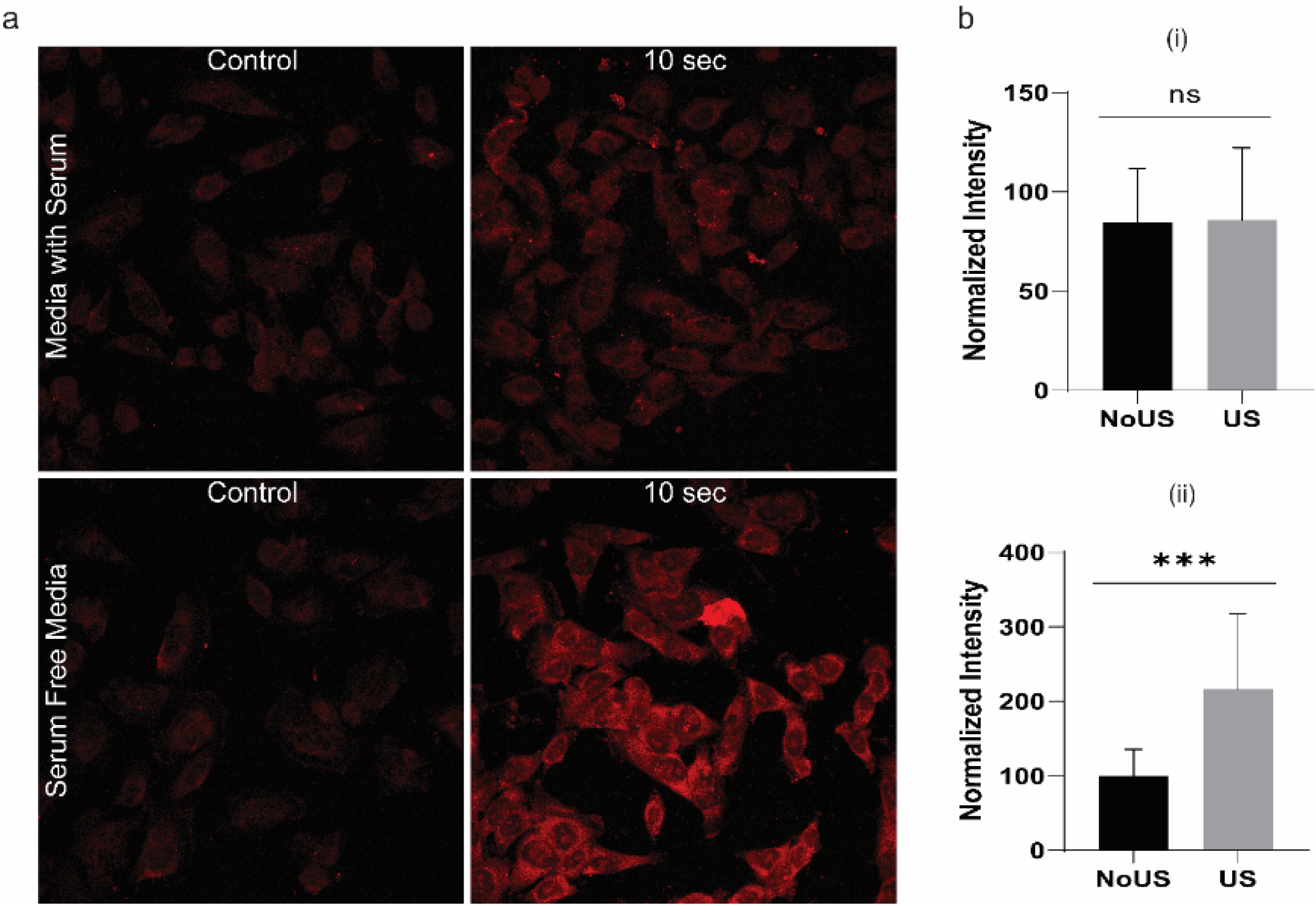
Effect of serum on 2µg/ml PI uptake. (a) Confocal images of cells showing the internalization of 2 µg/ml PI by SUM159 cells upon ultrasound exposure in the presence and absence of serum. Images were taken in Z-stacks and analyzed using FIJI ImageJ. (b) Quantification of PI permeation at 2 µg/ml concentration in (i) complete media (ii) serum-free media. The graph was plotted using GraphPad Prism 8. Error bar indicates mean ± SD of PI intensity in each condition. Normalized intensity was calculated from > 30 cells per condition. The complete media showed ns: non-significant difference in control and treated group whereas in serum free media *** indicates p value <0.005

We then investigated the effect of acoustic intensities ranging 2.8 W/cm^2^ to 8.4 W/cm^2^ in serum-free media with PI concentration of 2 µg/ml **(Figure 3a)**. We found that the penetration of PI was the highest at 5.6 and 7 W/cm^2^ and decreased at 8.4 W/cm^2^ **(Figure 3b)**. Further, we monitored the effect of time on PI’s penetration at constant US intensity of 5.6 W/cm^2^ **(Figure 3c)**. We tried different time points ranging from 10 sec to 120 sec and found that penetration was detected at 10 sec and increased up to 40 sec with the subsequent reduction afterward **(Figure 3d)**. The incubation time of PI after US treatment was kept constant in all the experiments as 10 mins. The controls were given no US treatment and incubated with PI for 10 mins. However, it was observed that with strong US exposure parameters (acoustic intensity and time), the morphology of the cells is altered. We thus chose an US exposure time of 10 sec for further experiments.

**Figure 3.**
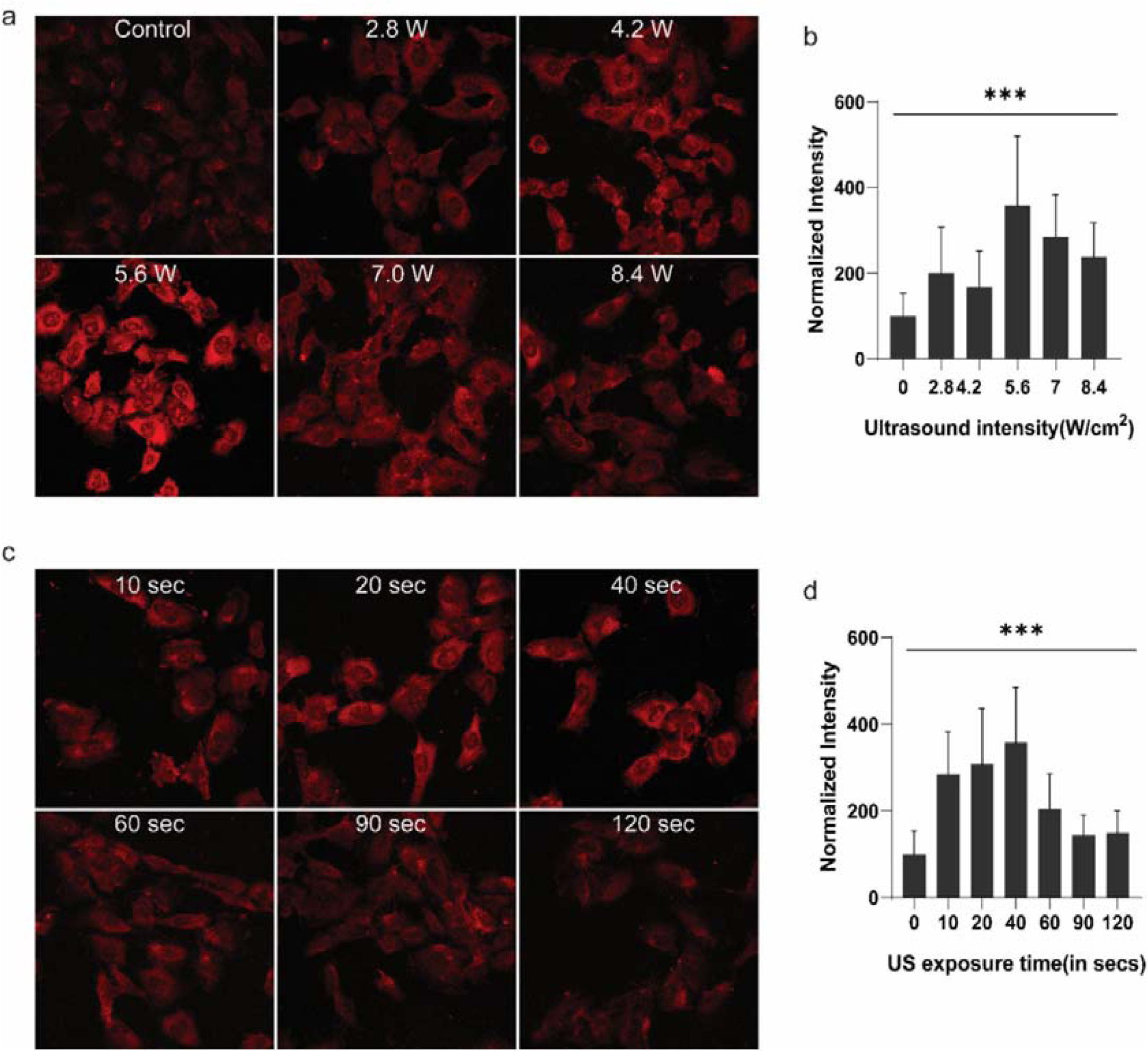
PI uptake in different acoustic settings in serum-free media. **(a)** Confocal images of intensity-dependent uptake of propidium iodide (PI) in breast cancer cells (SUM159) under ultrasound. Images were taken in Z-stacks and analyzed using FIJI ImageJ. **(b)** Quantification of PI penetration upon US treatment from cells in panel (a). The intensity of 5.6 and 7 W/cm^2^ showed the highest penetration. **(c)** Confocal images of cells showing time-dependent penetration of PI. **(d)** Quantification of PI penetration upon US treatment for different time points from cells in panel (c). The graph was plotted using GraphPad Prism 8. Error bar indicates mean ± SD of PI intensity in each condition. The normalized intensity was calculated from > 30cells per condition. *** indicates p-value <0.005

### 2. Characterization and cellular uptake of different molecular-weight fluorescent dextrans

Dynamic light scattering of different molecular weights (4, 10, 20, 40, 70, 150, 250, 500, 2000kDa) of FITC-Dextran’s was performed to analyze their hydrodynamic size. The graph shows that the average hydrodynamic radius (nm) increases with molecular weight **(Figure 4)**.

**Figure 4.**
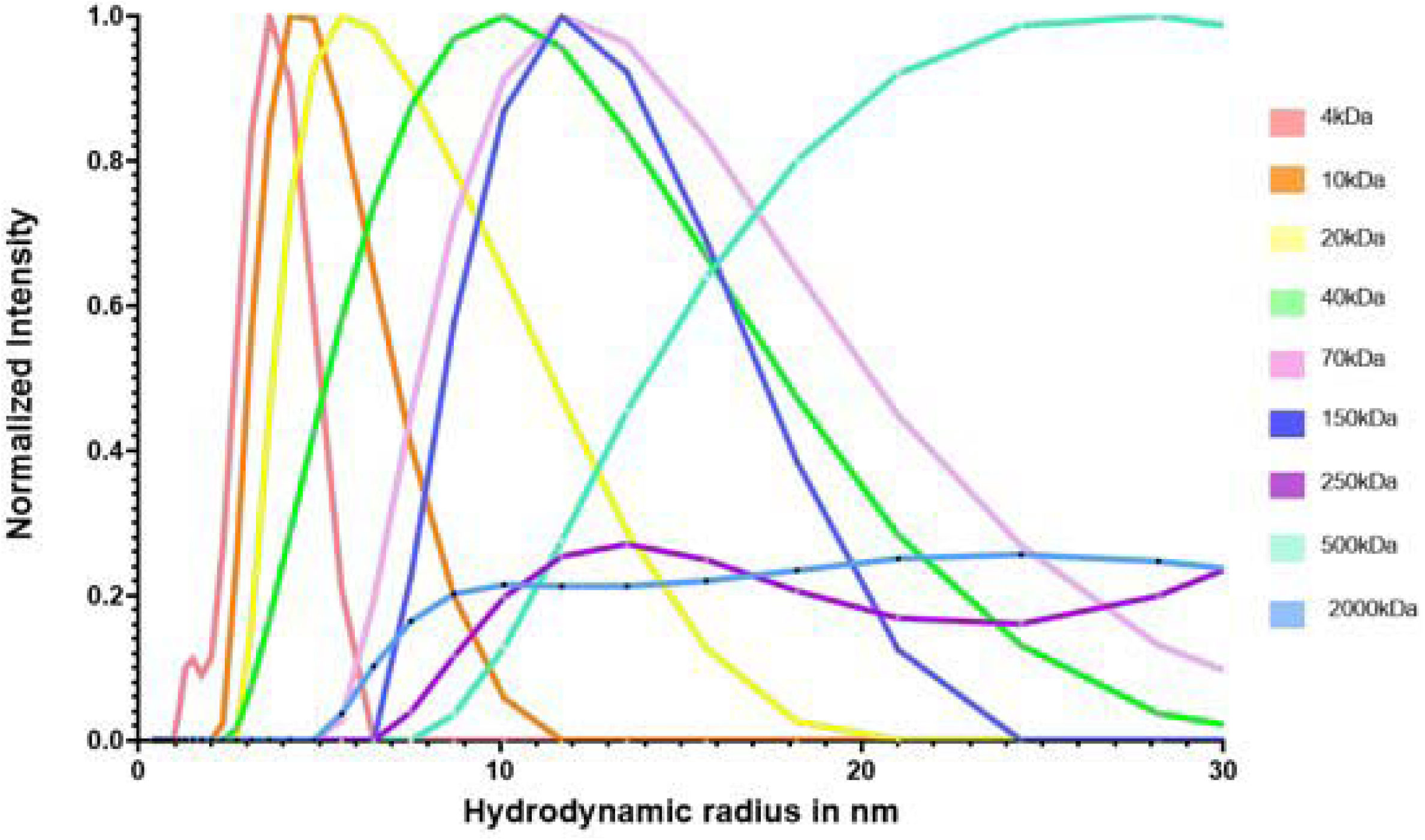
DLS of FITC-Dextran’s. Dynamic light scattering (DLS) of FITC-Dextran’s of different molecular weights (4, 10, 20, 40, 70, 150, 250, 500, 2000kDa).

FITC-Dextrans were been used as a marker to study fluid phase endocytosis in cells. We tested different acoustic intensities ranging from 2.4 to 8.4 W/cm2 for 10 s followed by incubation with (1 mg/ml) 4kDa FITC-dextran for 10 mins in serum-free media **(Figure 5a)**. The controls cells were given no US treatment and only incubated with FITC-dextran for 10 mins. The permeability of FITC-dextran increases with the increase in acoustic intensity for upto 5.6 W/cm2 and then decreases with higher intensity **(Figure 5b)**.

**Figure 5.**
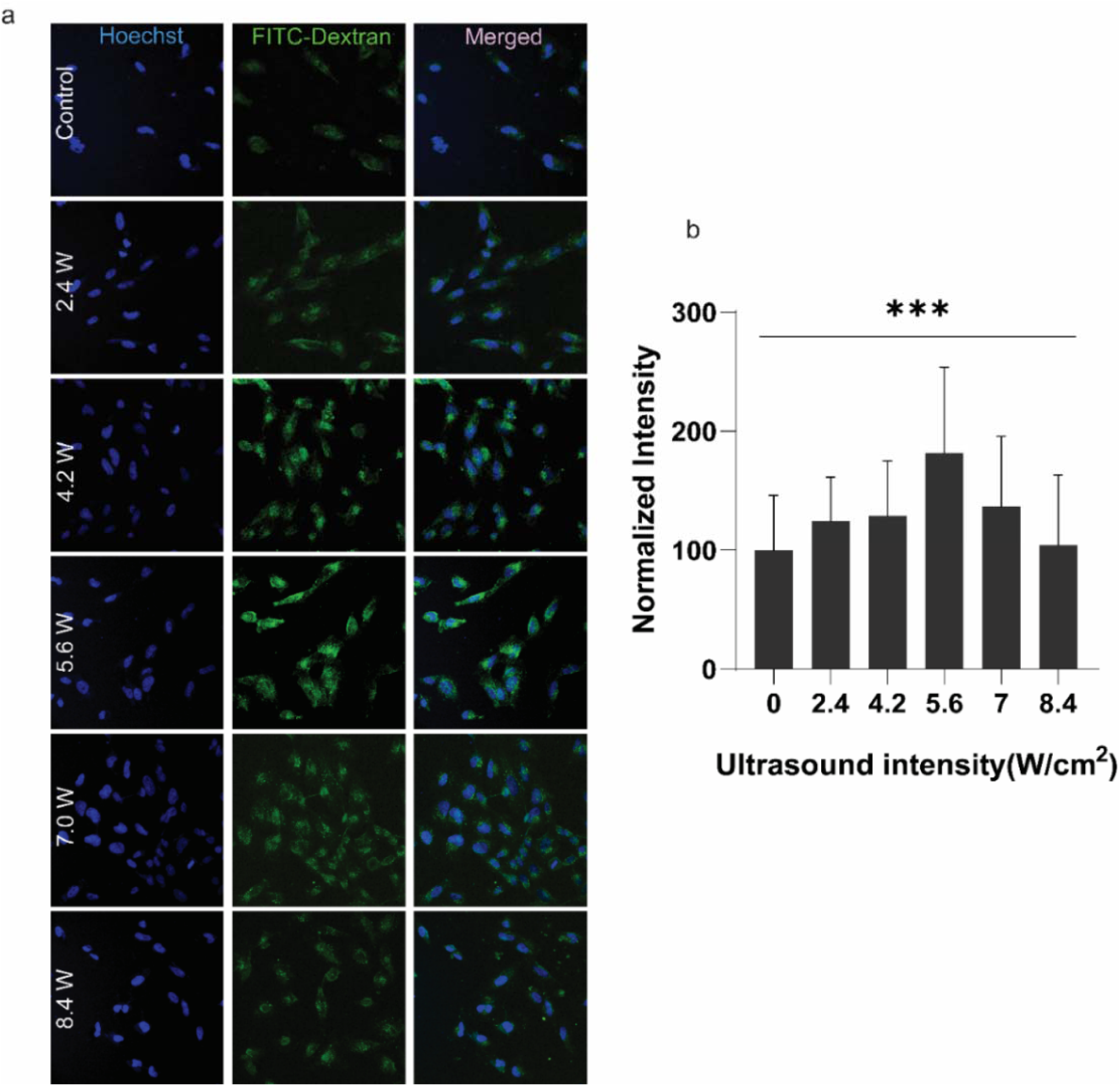
Optimization of US parameters for FITC-Dextrans. **(a)** Confocal images of SUM159 cells showing permeabilization of 4 kDa FITC-dextran. The fluorescence intensity was maximum in 5.6 W/cm^2^ **(b)** Quantification of FITC-dextran permeabilization upon US treatment with different intensities from cells in panel **(c)**. Error bar indicates mean ± SD of FITC-dextran intensity in each condition. The normalized intensity was calculated from > 30 cells per condition. *** indicates p-value <0.005.

We further checked the effect of molecular weight in the uptake of FITC-dextran upon US treatment. We used FITC-dextran particles with molecular weight ranging from 4 kDa to 2000 kDa in SUM159 cells. The cells were exposed to a constant US intensity of 5.6 W/cm^2^ for 10 s followed by incubation with 1 mg/ml FITC-dextran for 10 mins. The control cells were not exposed to US treatment and were incubated with FITC dextran under similar condition. The fluorescence intensity increased in the treated cells compared to non-treated cells. However, in 20 kDa, 40 kDa and 250 kD particles, there was no significant increase in the uptake upon US treatment (**Figure 6**).

**Figure 6.**
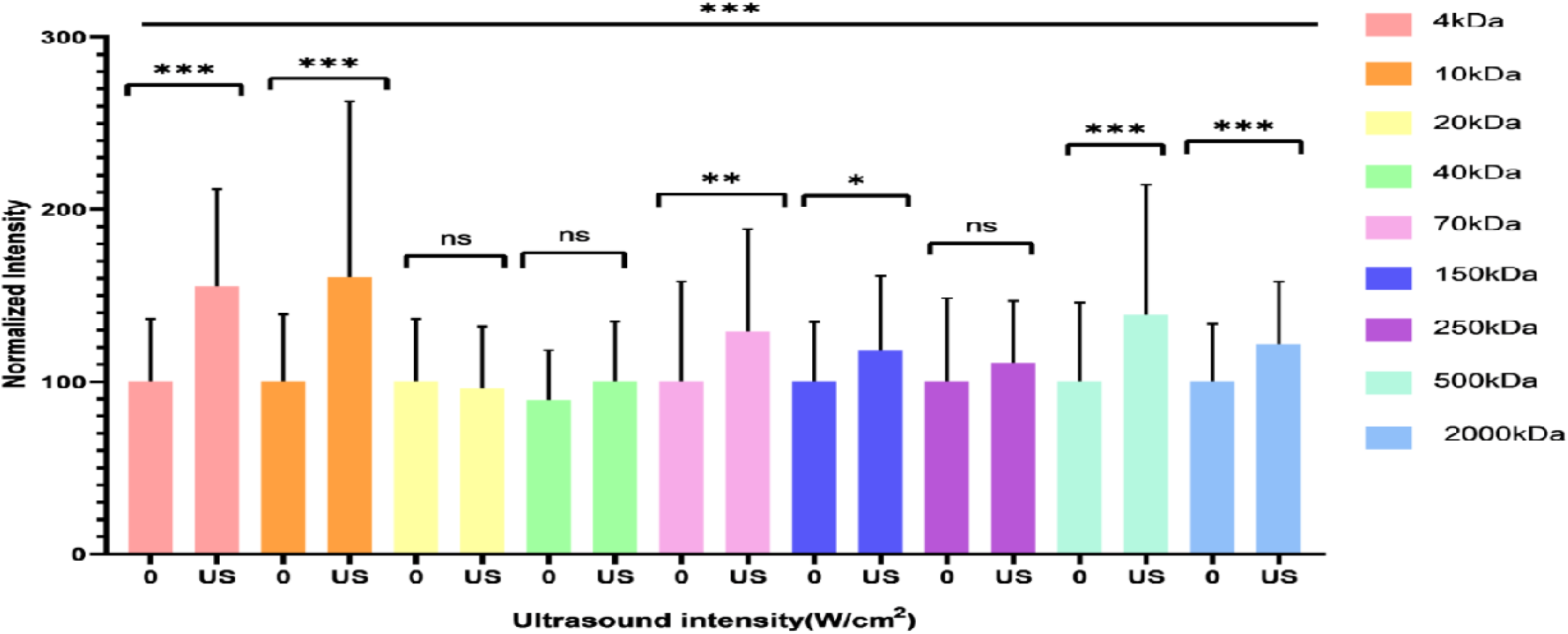
Effect of US treatment and molecular weight in internalization of FITC-dextran in SUM159 cells. Cellular fluorescence intensities of different molecular weights of FITC-Dextran’s (4, 10, 20, 40, 70, 150, 250, 500, 2000 kDa) in SUM159 cells in without or with US exposure shows enhancement of cellular fluorescence intensities of uptake FITX dextrans upon US exposure. *** = p < 0.0001, **=p=0.0075, *=p-=0.0136 (p<0.0001 in one-way ANOVA test).

### 3. Synthesis, characterization and cellular uptake of DNA tetrahedron

DNA tetrahedron (TD) has shown tremendous applications in biosensing and biomedical applications. DNA tetrahedron was synthesized using four single-stranded oligonucleotides through single-step assembly approach **(Figure 7a)**. In this method, the equimolar ratio of all the oligonucleotides is mixed and heated at a higher temperature of 95°C and gradually cooled to 4°C. We have synthesized two different sizes of DNA TD, i.e., small (TDS) of size ∼10nm and large (TDL) size ∼28nm. The preliminary characterization was done using electrophoretic mobility shift assay in which the mobility of higher-order structure is retarted. The ladder-like band in gel electrophoresis confirms the formation of a higher-order structure **(Figure 7b)**.

**Figure 7.**
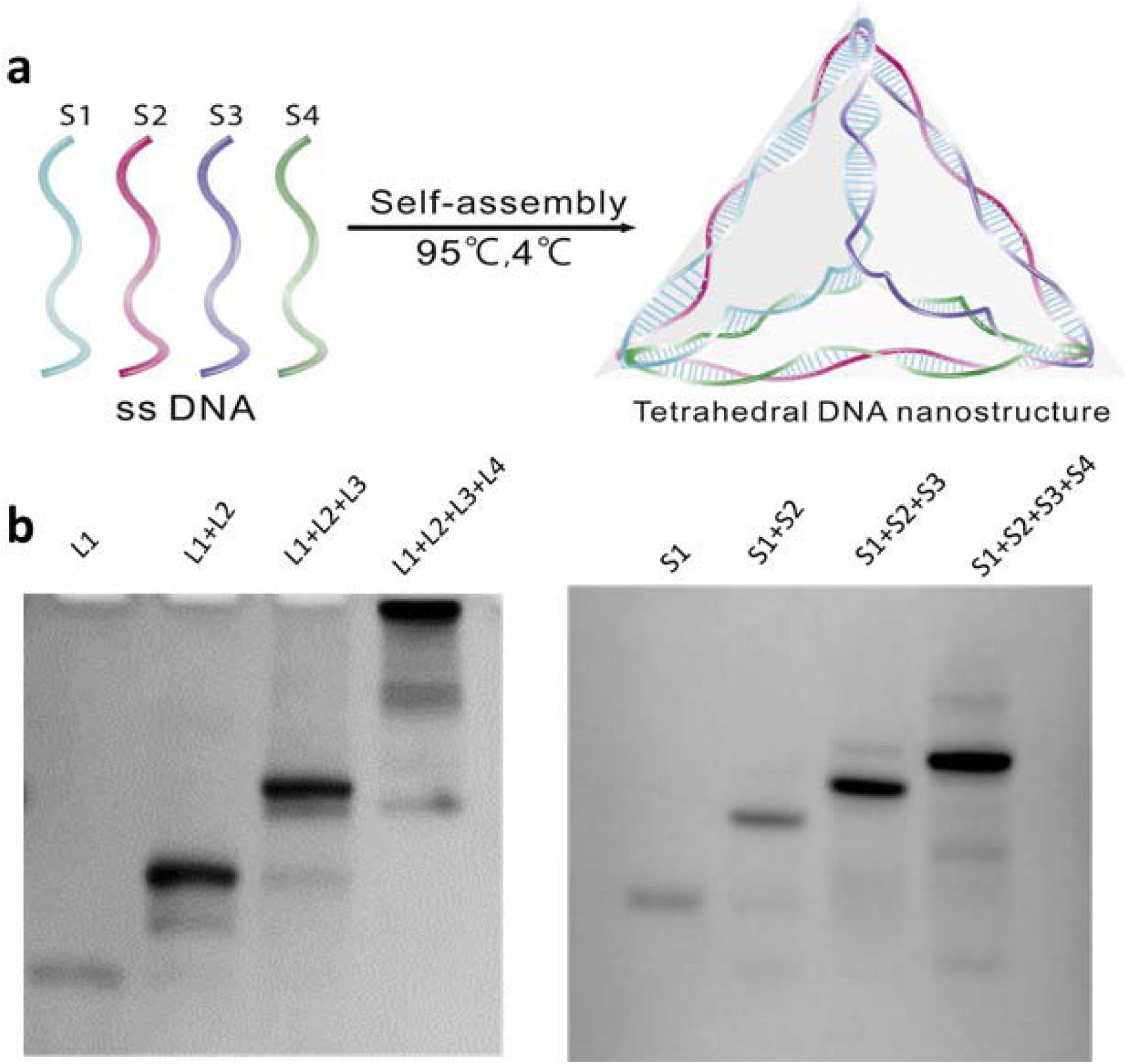
Synthesis and characterization of DNA tetrahedron. **(a)** One-pot synthesis of DNA tetrahedron [63]. **(b)** Gel-based characterization to confirm the formation of DNA TDL (larger TDN) and TDS (smaller TDN).

We further investigated the effect of acoustic intensities in the internalization of DNA TD. The internalization of TD was traced using Cy3 labeled oligonucleotide, which was incorporated during the assembly process. The uptake of both small and large DNA tetrahedrons (TDS and TDL) was observed under the influence of ultrasound with varying intensities ranging from 4.2 to 8.4 W/cm^2^ **(Figure 8, 9)**. The cells were exposed to the US for 10 s followed by incubation with 100 nM of DNA TD for 10 min. The controls were given no US exposure and only incubated with DNA TD for 10 min **(Figure 8a, 9a)**. The internalization of TD was increased upon US treatment, and for TDS, the internalization was maximum at 8.4 and 4.2 W/cm^2^ **(Figure 8b)**. However, for TDL, the internalization was maximum at 4.2, followed by 7 and 8.4 W/cm^2^ **(Figure 9b)**.

**Figure 8.**
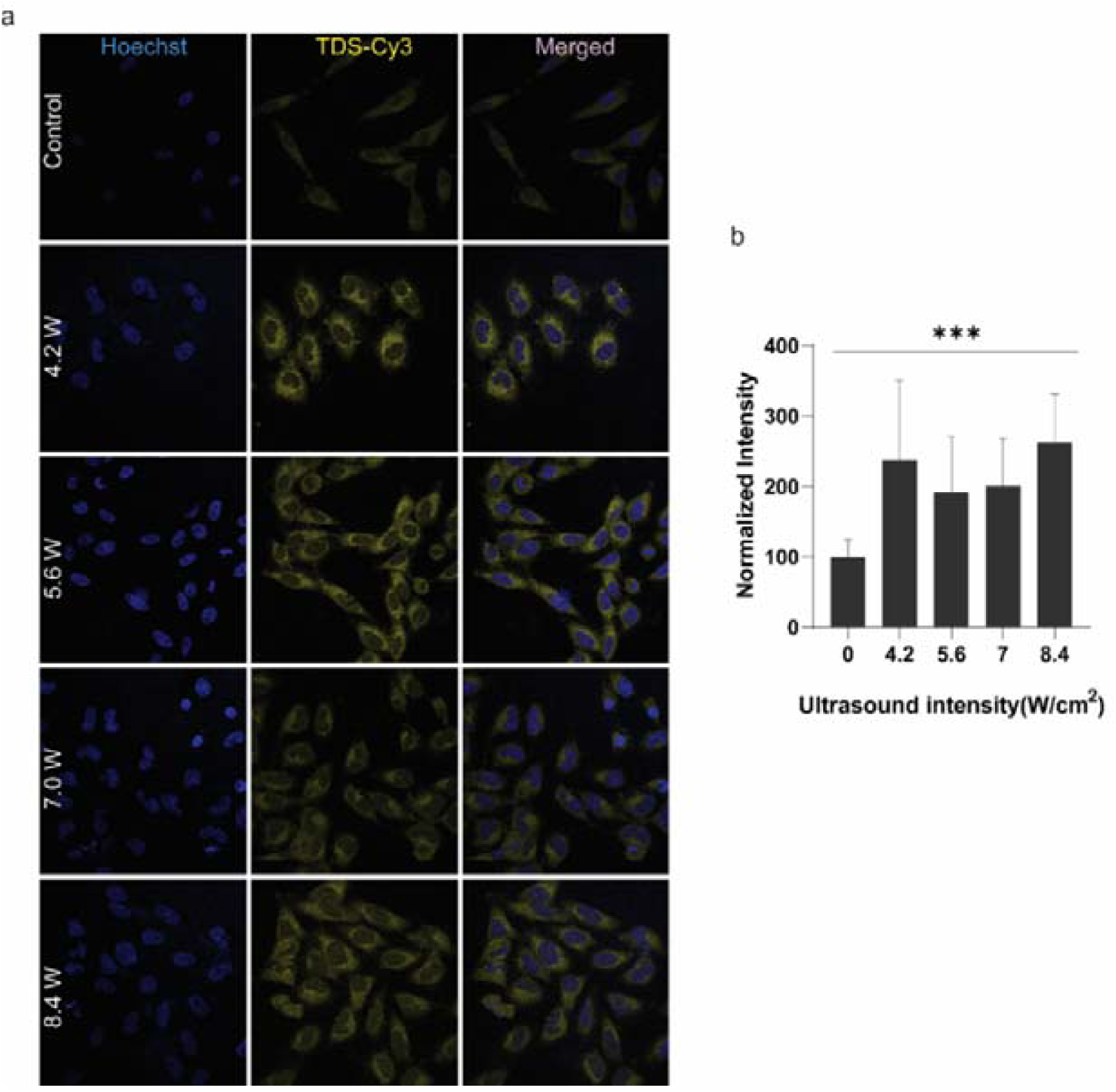
Cellular uptake of TDS in SUM159 cells upon US treatment. **(a)** Confocal images of cells showing the internalization of DNA TD upon US treatment with different intensities. Images were taken in Z-stacks and analyzed using FIJI ImageJ. **(b)** Quantification of TDS internalization upon US treatment from cells in panel (a). Error bar indicates mean ± SD of TDS intensity in each condition. The normalized intensity was calculated from > 30 cells per condition. *** indicates p <0.005

**Figure 9.**
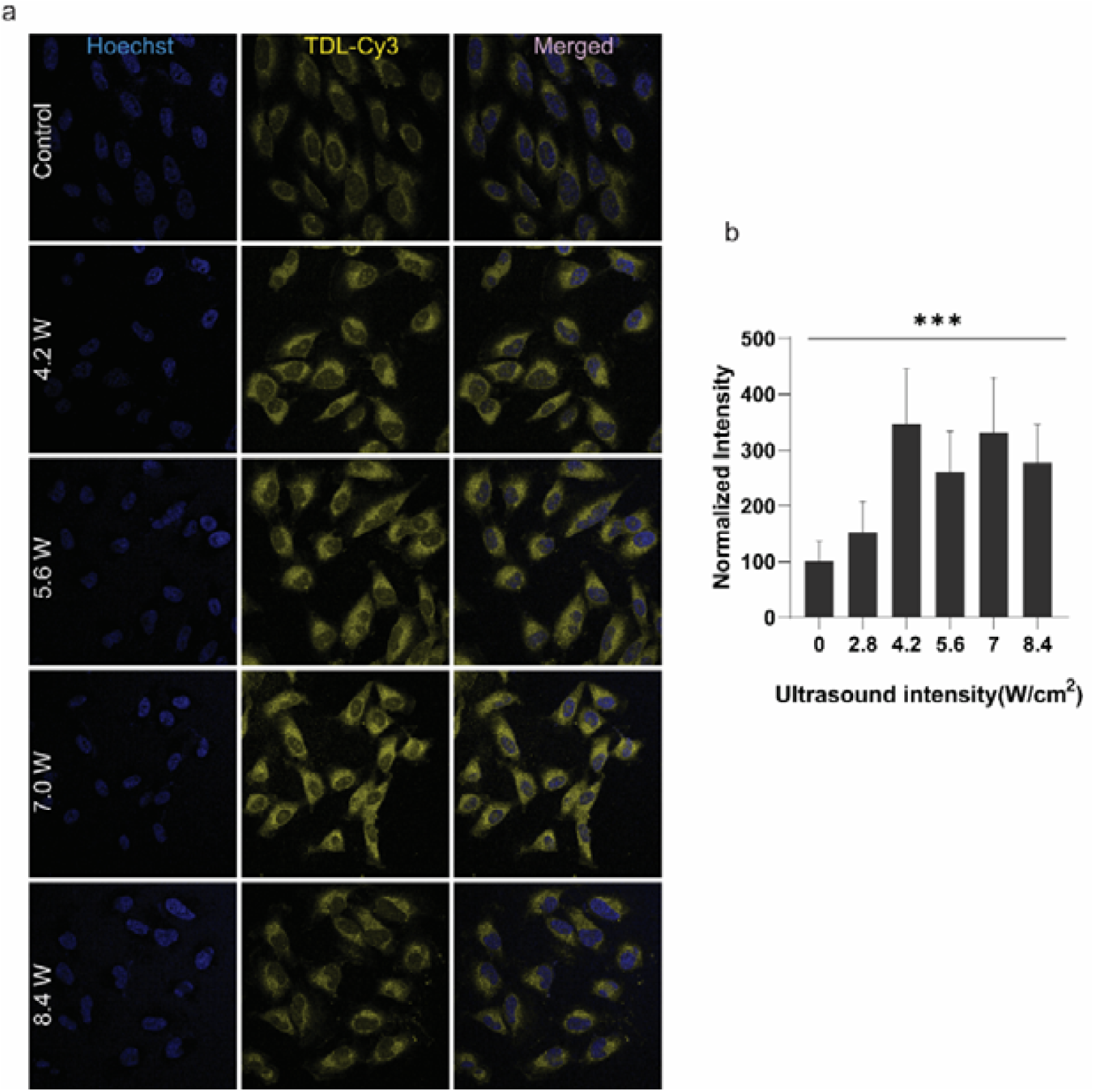
Cellular uptake of TDL in SUM159 cells upon US treatment. (a) Confocal images of cells showing the internalization of DNA TDL upon US treatment with different intensities. (b) Quantification of TDL internalization upon US treatment from cells in panel (a). The graph was plotted using GraphPad Prism 8. Error bar indicates mean ± SD of TDL intensity in each condition. The normalized intensity was calculated from > 30 cells per condition. *** indicates p <0.005

## Discussion

Acoustic cavitation mostly affects the cells by three mechanisms: making pores in the cell membrane (sonoporation), increasing endocytosis (stimulates active uptake), by opening cell-to-cell contacts. Propidium iodide is a DNA binding, cellular impermeable dye that can’t cross the cell membrane of a viable cell. Hence, it is used for staining necrotic or apoptotic cells [Yang, Y., Xiang, 2016, Tom van Rooij, 2016]. However, it can enter a membrane-compromised cell due to sonoporation through the pores formed in the membrane and acts as a marker of cell permeation or sonoporation tracer. A significant increase of the fluorescence in cells indicates sonoporation, as seen from the results of this study.

Dextrans are neutral macromolecules that are hydrophilic and non-toxic to humans [Lénárt P, 2006]. They are available in many molecular sizes they are often used as a molecular size marker for studying cell permeability [J.N. BeMiller, 2003]. To investigate the potential effects of molecular weight on ultrasound-mediated cellular uptake, we used sizes ranging from 4 kDa to 2000kDa [Afadzi M, Strand SP, 2013]. It’s been visible that cellular fluorescent intensity increases under the effect of ultrasound in almost all the FITC-Dextran’s. Also, a visible decreasing trend also observed while comparing different molecular weight dextrans. This trend seems consistent with the results seen in the research in context to the FITC-dextran’s endocytosis by ultrasound. The mechanism of acoustic cellular uptake is related to pore formation as well as enhanced endocytosis. It’s been seen that amount of internalized dextrans in small sizes dextrans can be significantly more although the FITC-label or fluorescent yield of 2000 kDa is high [M. Afadzi et al., 2013]. However, the number of vesicles present in the sonoporated cells can be more in high molecular dextrans [J. Hauser, 2009]. Meijering et al. reported that larger dextran’s (500 kDa) are taken up mainly through endocytosis and not through pore formation, whereas the smaller dextran’s (4 kDa) are taken up through both endocytosis and pores.

DNA nanostructures have high DNA density and due to which it is not easy to cross the membrane barrier. We choose DNA tetrahedron consists of four single-chain DNA strands which self-assemble by base pairing. In theory, DNA is confronted with membranes barriers due to its abundant charges. Some researchers presume that they interact with scavenger receptors or TLRs, leading to harmful signaling cascades when crossing the membrane [Hu, Y., Chen, 2017]. US-mediated uptake for DNA tetrahedron inside the cells could help overcome this barrier. By using ultrasound, we observed increased uptake of DNA tetrahedron for both sizes-small and large, indicating the active remodulation of plasma membrane by ultrasound thereby mediating its cellular uptake. The mechanisms of the uptake using ultrasound remain to be explored for these types of cargo.

## Conclusions

We demonstrated ultrasound-enhanced delivery of DNA tetrahedrons into breast cancer cells in an in vitro model. The favorable parameters for US intensity exposure time were evaluated *in vitro*. An increase in cellular uptake of membrane impermeable dye Propidium iodide (PI) with respect to the controls at the ultrasound intensities employed in this study suggests that sonoporation was the mechanism for enhanced delivery of DNA nanocages. At the exposure intensities applied, no visible effects were observed in the morphology of the cells post treatment. Further, FITC-Dextrans were tested for their uptake in SUM159 cells. Working ultrasound parameters for dextrans were found in the optimization process. Enhanced uptake of various molecular weights dextrans further suggests that sonoporation was induced upon exposure to ultrasound. There was a trend observed in the uptake of different molecular weights of FITC-dextran. Our initial results have setup the groundwork for ultrasound-enhanced delivery of DNA based agents as well as other nanoscale cargo without using any external cavitation nuclei or contrast agents; which can have multiple biological and biomedical applications.

## Materials and methods

### Cell culture

Human breast cancer cells (SUM159) and human retinal eye pigment epithelial (RPE-1) cells was cultured in DMEM (RPE-1) and HamsF12 (for SUM159) media respectively with 10% FBS in 30 °C, 5% CO2 and 90% humidity. Cells were seeded in well plates for the subsequent experiment as per the need of number of cells in experiment.

### Ultrasound treatment

The therapeutic ultrasound machine used was dual channel therapy transducer Lcs-128. The ultrasound apparatus consists of the ultrasound machine, ultrasound probe holder, ultrasound transducer and ultrasound jelly. First, a probe holder was used to hold the US probe such that the transducer surface was in the vertical direction facing upward. Then, US jelly was applied. The culture dish containing coverslip and media was kept on it and US treatment was given to the cells. The ultrasound parameters used were as follows-US frequency -1MHz, Pulse repetition frequency (PRF) - 10Hz, Duty Cycle (DC)-5%. The US intensity and US exposure time was optimized later in the experiments.

### PI uptake in SUM159 cells

Propidium iodide (PI) is a red-fluorescent nuclear and chromosome counterstain. As, it is impermeable to the cell membrane, the formation of pores after US treatment can make it permeable to the cells. For the experiment, SUM159 cells were plated on 18 mm coverslips and incubated at 37°C. Working solution of 1 µg/ml Propidium iodide (PI) in HamsF12 complete media was prepared. In one set of experiment, the US exposure time was kept constant i.e.,10 sec. US intensity was varied between the values-2.8, 4.2, 5.6, 7, 8.4 W/cm^2^. In another set of experiment, the US intensity was kept constant i.e.,4.2 W/cm^2^. US exposure time was varied between the values-10, 20, 40, 60, 120 s. immediately before the US treatment, 1 ml of the PI working solution was added in the 35 mm culture dish containing the coverslip. After US treatment, the cells were incubated for 10 min. Controls were given no US treatment. Cells were washed thrice by PBS and fixed with 4% PFA. Coverslips were prepared and imaging was performed using confocal microscopy. Experiments were also repeated with the HamsF12 serum free media.

### DLS of FITC-Dextrans

1 mg/ml working solutions of different molecular weights of Fluorescein isothiocyanate Dextran’s (4, 10, 20, 40, 70, 150, 250, 500, 2000 kDa) were prepared using filtered distilled water. Dynamic Light Scattering (DLS) was performed to analyze the prepared solutions. The size-distribution profiles of different molecular weights of FITC-Dextran’s were determined and compared.

### Endocytosis of FITC-Dextrans in SUM159 cells

This experiment was done to optimize US intensity and US exposure time for uptake of FITC-Dextran’s. SUM159 cells were plated on 18 mm coverslips and incubated at 37°C for the experiment. Working solution of 1 mg/ml Fluorescein isothiocyanate Dextran (4 kDa) in HamsF12 serum free media was prepared. In one set of experiment, the US exposure time was kept constant i.e.,10 sec. US intensity was varied between the values-2.8, 4.2, 5.6, 7, 8.4 W/cm^2^. In another set of experiment, the US intensity was kept constant i.e., 4.2 W/cm^2^. US exposure time was varied between the values-10, 20, 40, 60, 120 s. immediately before the US treatment, 1 ml of the working solution was added in the 35 mm culture dish containing the coverslip. After US treatment, the cells were incubated for 15 min. No US treatment was given to the controls. Cells were washed thrice by PBS. Coverslips were prepared and imaging was perfomed using confocal microscopy. Further, the optimized US intensity and US exposure time parameters for 4 kDa FITC-Dextran were used to study the uptake of different molecular weights of Fluorescein isothiocyanate Dextran’s (4, 10, 20, 40, 70, 150, 250, 500, 2000 kDa) in SUM159 cells by using the same protocol.

### DNA tetrahedron synthesis

One pot synthesis of DNA tetrahedron of different sizes (small (8-10 nm) and large (28-30 nm)) was performed by Polymerase chain reaction (PCR). Four small DNA oligomers were used to make a tetrahedron. For every 20 µL of solution, 5 µL of each oligomer was added from a working stock (10 µM) kept at 20° C. In addition, to make the final structure stable, 2 mM of MgCl_2_ was added in the same solution. The remaining volume was filled with 1X PBS. The PCR tubes containing the solution were run for one PCR cycle of 4.5 h. In this PCR cycle, the temperature was increased to 95°C followed by a 5°C decrease every 15 minutes. At the final temperature of 4°C, the cycle tops.

### Native Polyacrylamide gel electrophoresis

To characterize the purity of DNA tetrahedron particles, four different solutions were prepared. Three of them were controls, containing only one strand, two strands and three oligomer strands. Other solution contained all the four strands thus, can form a DNA tetrahedron. 10% Native-PAGE was performed for all four solutions. The gel and running buffer were 1xTAE buffer. 4µL samples were loaded with 1.5 µL purple/orange tracking dye and 0.4 µL gel buffer. The electrophoresis was done at 60 V for 80 minutes.

### Uptake of DNA tetrahedron in SUM159 cells

This experiment was done to optimize ultrasound parameters-US intensity and US exposure time for DNA tetrahedron. SUM159 cells were plated on 18 mm coverslips and incubated at 37°C for the experiment. Working solution of DNA tetrahedron (small - 83.3 nM and large - 100nM) was prepared in HamsF12 serum free media. In one set of experiment, the US exposure time was kept constant i.e.,10 sec. US intensity was varied between the values-2.8, 4.2, 5.6, 7, 8.4 W/cm^2^. In another set of experiment, the US intensity was kept constant i.e., 4.2 W/cm^2^. US exposure time was varied between the values-10, 20, 40, 60, 120 sec. Immediately before the US treatment, 1 ml of the working solution was added in the 35 mm culture dish containing the coverslip. After US treatment, the cells were incubated for 10 min. Controls were given no US treatment. Cells were washed thrice by PBS. Cells were fixed with 4% PFA. Coverslips were prepared and imaging was done in confocal microscopy. The whole experiment was performed with the both small as well as large sizes of DNA tetrahedrons.

### Image analysis and statistics

All confocal images were taken in Z-stack and compiled together using FIJI ImageJ (https://imagej.net/Fiji). The cells were marked and intensities were collected and stored in excel sheets. To check the significance of the data, GraphPad Prism 8 was used. Student t-test and One-way ANOVA tests were used to test statistical significance. P-value less than 0.05 was considered statistically significant.

## Acknowledgments

We sincerely thank the members of DB and HS groups for critically reading the manuscript and their valuable feedback. KB and AR thank IITGN-MHRD, GoI for the fellowship. DB and HS thank SERB, GoI for Ramanujan Fellowship and Startup grant, DB, KMS, HS thank IITGN for the startup funds and Department of Biotechnology GoI for Har Govind Khorana Innovative Young Biotechnologist Award. DB thanks BRNS-DAE for the research grant. KB and CKJ thank IITGN-MHRD for MTech and Postdoctoral fellowships respectively. The works in the host labs are also funded by GSBTM, Govt of Gujarat, India.

The authors declare no conflict of interest.

## Notes

### Competing Interest Statement

The authors have declared no competing interest.

## References

E. N. Harvey, “BIOLOGICAL ASPECTS OF ULTRASONIC WAVES, A GENERAL SURVEY,” The Biological Bulletin, vol. 59, no. 3, pp. 306–325, Dec. 1930, doi: 10.2307/1536819.

Z. Izadifar, P. Babyn, and D. Chapman, “Mechanical and Biological Effects of Ultrasound: A Review of Present Knowledge,” Ultrasound in Medicine and Biology, vol. 43, no. 6. Elsevier USA, pp. 1085–1104, Jun. 01, 2017, doi: 10.1016/j.ultrasmedbio.2017.01.023.

I. Lentacker, I. de Cock, R. Deckers, S. C. de Smedt, and C. T. W. Moonen, “Understanding ultrasound induced sonoporation: Definitions and underlying mechanisms,” Advanced Drug Delivery Reviews, vol. 72. Elsevier, pp. 49–64, Jun. 15, 2014, doi: 10.1016/j.addr.2013.11.008.

H. Chen and E. E. Konofagou, “The size of blood-brain barrier opening induced by focused ultrasound is dictated by the acoustic pressure,” Journal of Cerebral Blood Flow and Metabolism, vol. 34, no. 7, pp. 1197–1204, 2014, doi: 10.1038/jcbfm.2014.71.

K. Kooiman et al., “Ultrasound-Responsive Cavitation Nuclei for Therapy and Drug Delivery,” Ultrasound in Medicine and Biology, vol. 46, no. 6. Elsevier USA, pp. 1296– 1325, Jun. 01, 2020, doi: 10.1016/j.ultrasmedbio.2020.01.002.

O. Couture, J. Foley, N. F. Kassell, B. Larrat, and J.-F. Aubry, “Review of ultrasound mediated drug delivery for cancer treatment: updates from pre-clinical studies,” Translational Cancer Research, vol. 3, no. 5, 2014, doi: 10.21037/3354.

Karimi M, Zangabad PS, Mehdizadeh F, Malekzad H, Ghasemi A, Bahrami S, Zare H, Moghoofei M, Hekmatmanesh A, Hamblin MR. Nanocaged platforms: modification, drug delivery and nanotoxicity. Opening synthetic cages to release the tiger. Nanoscale. 2017 Jan 26;9(4):1356–1392. doi: 10.1039/c6nr07315h. PMID: 28067384; PMCID: PMC5300024.

Fan, Z., Kumon, R. E., Park, J., & Deng, C. X. (2010). Intracellular delivery and calcium transients generated in sonoporation facilitated by microbubbles. Journal of controlled release:official journal of the Controlled Release Society, 142(1), 31– 39. https://doi.org/10.1016/j.jconrel.2009.09.031

Li, F., Yang, C., Yuan, F., Liao, D., Li, T., Guilak, F., & Zhong, P. (2018). Dynamics and mechanisms of intracellular calcium waves elicited by tandem bubble-induced jetting flow. Proceedings of the National Academy of Sciences of the United States of America, 115(3), E353–E362. https://doi.org/10.1073/pnas.1713905115

Wang, M., Zhang, Y., Cai, C. et al. Sonoporation-induced cell membrane permeabilization and cytoskeleton disassembly at varied acoustic and microbubble-cell parameters. Sci Rep 8, 3885 (2018). https://doi.org/10.1038/s41598-018-22056-8

Duan, X., Zhou, Q., Wan, J.M.F. et al. sonoporation generates downstream cellular impact after membrane resealing. Sci Rep 11, 5161 (2021). https://doi.org/10.1038/s41598-021-84341-3,

Chen Xian, Leow Ruen Shan, Hu Yaxin, Wan Jennifer M.F. and Yu Alfred C.H. 2014 Single-site sonoporation disrupts actin cytoskeleton organizationJ. R. Soc. Interface.112014007120140071 https://doi.org/10.1098/rsif.2014.0071

Yang, Y., Xiang, Y. & Xu, M. From red to green: the propidium iodide-permeable membrane of Shewanella decolorationis S12 is repairable. Sci Rep 5, 18583 (2016). https://doi.org/10.1038/srep18583

Kiselevsky, D.B., Samuilov, V.D. Permeability of the Plasma Membrane for Propidium Iodide and Destruction of Cell Nuclei in the Epidermis of Pea Leaves: The Effect of Polyelectrolytes and Detergents. Moscow Univ. Biol.Sci. Bull. 74, 147–153 (2019). https://doi.org/10.3103/S0096392519030052

R. Karshafian and P. N. Burns, “Ultrasound and microbubble mediated generation of transient pores on cell membranes in vitro,” 2009 IEEE International Ultrasonics Symposium, 2009, pp. 1805–1808, doi: 10.1109/ULTSYM.2009.5441643.

Yoon, S., Kim, M. G., Chiu, C. T., Hwang, J. Y., Kim, H. H., Wang, Y., & Shung, K. K. (2016). Direct and sustained intracellular delivery of exogenous molecules using acoustic-transfection with high frequency ultrasound. Scientific reports, 6, 20477. https://doi.org/10.1038/srep20477

Ng KY, Liu Y. Therapeutic ultrasound: its application in drug delivery. Med Res Rev. 2002 Mar;22(2):204–23. doi: 10.1002/med.10004. PMID: 11857639.

Tom van Rooij, Ilya Skachkov, Inés Beekers, Kirby R. Lattwein, Jason D. Voorneveld, Tom J.A. Kokhuis, Deep Bera, Ying Luan, Antonius F.W. van der Steen, Nico de Jong, Klazina Kooiman, Viability of endothelial cells after ultrasound-mediated sonoporation: Influence of targeting, oscillation, and displacement of microbubbles, Journal of Controlled Release, Volume 238, 2016,Pages 197–211,ISSN 0168-3659,https://doi.org/10.1016/j.jconrel.2016.07.037.

Afadzi M, Strand SP, Nilssen EA, Måsøy SE, Johansen TF, Hansen R, Angelsen BA, de L Davies C. Mechanisms of the ultrasound-mediated intracellular delivery of liposomes and dextrans. IEEE Trans Ultrason Ferroelectr Freq Control. 2013 Jan;60(1):21–33. doi: 10.1109/TUFFC.2013.2534. PMID: 23287910.

Wilms CD, Häusser M. Twitching towards the ideal calcium sensor. Nat Methods. 2014 Feb;11(2):139–40. doi: 10.1038/nmeth.2814. PMID: 24481218.

Zhou Y, Shi J, Cui J, Deng CX. Effects of extracellular calcium on cell membrane resealing in sonoporation. J Control Release. 2008 Feb 18;126(1):34–43. doi: 10.1016/j.jconrel.2007.11.007. Epub 2007 Nov 22. PMID: 18158198; PMCID: PMC2270413.

Greenleaf WJ, Bolander ME, Sarkar G, Goldring MB, Greenleaf JF. Artificial cavitation nuclei significantly enhance acoustically induced cell transfection. Ultrasound Med Biol. 1998 May;24(4):587–95. doi: 10.1016/s0301-5629(98)00003-9. PMID: 9651968.

Hwang JY, Lim HG, Yoon CW, Lam KH, Yoon S, Lee C, Chiu CT, Kang BJ, Kim HH, Shung KK. Non-contact high-frequency ultrasound microbeam stimulation for studying mechanotransduction in human umbilical vein endothelial cells. Ultrasound Med Biol. 2014 Sep;40(9):2172–82. doi: 10.1016/j.ultrasmedbio.2014.03.018. Epub 2014 Jul 9. PMID: 25023109; PMCID: PMC4130794.

Lénárt P, Ellenberg J. Monitoring the permeability of the nuclear envelope during the cell cycle. Methods. 2006 Jan;38(1):17–24. doi: 10.1016/j.ymeth.2005.07.010. PMID: 16343937.

J.N. BeMiller, DEXTRAN,Editor(s): Benjamin Caballero, Encyclopedia of Food Sciences and Nutrition (Second Edition), Academic Press, 2003, Pages 1772–1773, ISBN 9780122270550,https://doi.org/10.1016/B0-12-227055-X/00330-8.

B. D. Meijering, L. J. Juffermans, A. van Wamel, R. H. Henning, I. S. Zuhorn, M. Emmer, A. M. Versteilen, W. J. Paulus, W. H. van Gilst, K. Kooiman, N. de Jong, R. J. Musters, L. E. Deelman, and O. Kamp, “Ultrasound and microbubble-targeted delivery of macromolecules is regulated by induction of endocytosis and pore formation,” Circ. Res., vol. 104, pp. 679–687, Mar. 13, 2009.

J. Hauser, M. Ellisman, H. U. Steinau, E. Stefan, M. Dudda, and M. Hauser, “Ultrasound enhanced endocytotic activity of human fibroblasts,” Ultrasound Med. Biol., vol. 35, pp. 2084–2092, Dec. 2009.

M. Afadzi et al., “Mechanisms of the ultrasound-mediated intracellular delivery of liposomes and dextrans,” in IEEE Transactions on Ultrasonics, Ferroelectrics, and Frequency Control, vol. 60, no. 1, pp. 21–33, January 2013, doi: 10.1109/TUFFC.2013.2534.

Khanna S, Hudson B, Pepper CJ, Amso NN, Coakley WT. Fluorescein isothiocynate-dextran uptake by chinese hamster ovary cells in a 1.5 MHz ultrasonic standing wave in the presence of contrast agent. Ultrasound Med Biol. 2006 Feb;32(2):289–95. doi: 10.1016/j.ultrasmedbio.2005.11.002. PMID: 16464674.

G. Apodaca, “Modulation of membrane traffic by mechanical stimuli,” Am. J. Physiol. Renal Physiol., vol. 282, pp. F179–F190, Feb. 2002

K. Lawler, G. O’Sullivan, A. Long, and D. Kenny, “Shear stress induces internalization of E-cadherin and invasiveness in metastatic oesophageal cancer cells by a Src-dependent pathway,” Cancer Sci., vol. 100, pp. 1082–1087, Jun. 2009.

Schwarz, D. S., & Blower, M. D. (2016). The endoplasmic reticulum: structure, function and response to cellular signaling. Cellular and molecular life sciences:CMLS, 73(1), 79–94. https://doi.org/10.1007/s00018-015-2052-6

Fan, Z., Kumon, R. E., & Deng, C. X. (2014). Mechanisms of microbubble-facilitated sonoporation for drug and gene delivery. Therapeutic delivery, 5(4), 467– 486. https://doi.org/10.4155/tde.14.10

Hu, Y., Chen, Z., Zhang, H., Li, M., Hou, Z., Luo, X., & Xue, X. (2017). Development of DNA tetrahedron-based drug delivery system. Drug delivery, 24(1), 1295– 1301. https://doi.org/10.1080/10717544.2017.1373166

Meijering BD, Juffermans LJ, van Wamel A, Henning RH, Zuhorn IS, Emmer M, Versteilen AM, Paulus WJ, van Gilst WH, Kooiman K, de Jong N, Musters RJ, Deelman LE, Kamp O. Ultrasound and microbubble-targeted delivery of macromolecules is regulated by induction of endocytosis and pore formation. Circ Res. 2009 Mar 13;104(5):679–87. doi: 10.1161/CIRCRESAHA.108.183806. Epub 2009 Jan 22. PMID: 19168443.

Walker, W. A., Tarannum, M., & Vivero-Escoto, J. L. (2016). Cellular Endocytosis and Trafficking of Cholera Toxin B-Modified Mesoporous Silica Nanoparticles. Journal of materials chemistry. B, 4(7), 1254–1262. https://doi.org/10.1039/C5TB02079D

Day, C. A., & Kenworthy, A. K. (2015). Functions of cholera toxin B-subunit as a raft cross-linker. Essays in biochemistry, 57, 135–145. https://doi.org/10.1042/bse0570135

Iglesias-Bartolomé R, Trenchi A, Comín R, Moyano AL, Nores GA, Daniotti JL. Differential endocytic trafficking of neuropathy-associated antibodies to GM1 ganglioside and cholera toxin in epithelial and neural cells. Biochim Biophys Acta. 2009 Dec;1788(12):2526–40. doi: 10.1016/j.bbamem.2009.09.018. Epub 2009 Oct 2. PMID: 19800863.

A. Carovac, F. Smajlovic, and D. Junuzovic, “Application of Ultrasound in Medicine,” Acta Informatica Medica, vol. 19, no. 3, p. 168, 2011, doi: 10.5455/aim.2011.19.168-171.

H. Kuttruff and H. Kuttruff, “Application of Ultrasound in Medical Diagnostics,” in Ultra sonics, Springer Netherlands, 1991, pp. 297–324.

E. N. Harvey, “BIOLOGICAL ASPECTS OF ULTRASONIC WAVES, A GENERAL SURVEY,” The Biological Bulletin, vol. 59, no. 3, pp. 306– 325, Dec. 1930, doi: 10.2307/1536819

R. Mundi, S. Petis, R. Kaloty, V. Shetty, and M. Bhandari, “Low-intensity pulsed ultrasound: Fracture healing,” in Indian Journal of Orthopaedics, Apr. 2009, vol. 43, no. 2, pp. 132–140, doi: 10.4103/0019-5413.50847.

P. D. McClain, J. N. Lange, and D. G. Assimos, “Optimizing shock wave lithotripsy: a comprehensive review.,” Reviews in urology, vol. 15, no. 2, pp. 49–60, 2013, Accessed: Jan. 28, 2021. [Online]. Available: http://www.ncbi.nlm.nih.gov/pubmed/24082843.

I. Lentacker, I. de Cock, R. Deckers, S. C. de Smedt, and C. T. W. Moonen, “Understanding ultrasound induced sonoporation: Definitions and underlying mechanism s,” Advanced Drug Delivery Reviews, vol. 72. Elsevier, pp. 49– 64, Jun. 15, 2014, doi: 10.1016/j.addr.2013.11.008.

Y. Hu, J. M. F. Wan, and A. C. H. Yu, “Membrane Perforation and Recovery Dynamics in Microbubble-Mediated Sonoporation,” Ultrasound in Medicine and Biology, vol. 39, no. 12, pp. 2393–2405, 2013, doi: 10.1016/j.ultrasmedbio.2013.08.003.

K. B. Bader and C. K. Holland, “Gauging the likelihood of stable cavitation from ultrasound contrast agents,” Physics in Medicine and Biology, vol. 58, no. 1, pp. 127– 144, Jan. 2013, doi: 10.1088/0031-9155/58/1/127.

Z. Izadifar, P. Babyn, and D. Chapman, “Mechanical and Biological Effects of Ultrasound: A Review of Present Knowledge,” Ultrasound in Medicine and Biology, vol. 43, no. 6. Elseviers USA, pp. 1085– 1104, Jun. 01, 2017, doi: 10.1016/j.ultrasmedbio.2017.01.023.

K. Kooiman et al., “Ultrasound-Responsive Cavitation Nuclei for Therapy and Drug Delivery,” Ultrasound in Medicine and Biology, vol. 46, no. 6. Elsevier USA, pp. 1296–1325, Jun. 01, 2020, doi: 10.1016/j.ultrasmedbio.2020.01.002.

Bhatia, D.; Arumugam, S.; Nasilowski, M.; Joshi, H.; Wunder, C.; Chambon, V.; Prakash, V.; Grazon, C.; Nadal, B.; Maiti, P. K.; Johannes, L.; Dubertret, B.; Krishnan, Y. Quantum Dot-Loaded Monofunctionalized DNA Icosahedra for Single-Particle Tracking of Endocytic Pathways. Nature Nanotechnology 2016, 11 (12),1112–1119. doi: 10.1038/nnano.2016.150.

Rajwar, A.; Kharbanda, S.; Chandrasekaran, A. R.; Gupta, S.; Bhatia, D. Designer, Programmable 3D DNA Nanodevices to Probe Biological Systems. ACS Appl. Bio Mater. 2020, 3 (11), 7265–7277. doi: 10.1021/acsabm.0c00916.

